# Beyond fish in formation: A two-tier approach for biomechanical studies of collective movement

**DOI:** 10.64898/2026.02.28.708741

**Authors:** Yangfan Zhang, Divya Ramesh, George V. Lauder

## Abstract

Despite much of the literature perceiving fish schooling as an organized system with a focus on fixed formations for theoretical analyses, experimental observations suggest that frequent positional rearrangement commonly occurs. Previous studies have also demonstrated that fish schools reduce locomotor costs relative to individuals swimming alone. This introduces an intriguing dichotomy. How can individual fish within schools exhibit dynamic interactions while also saving energy? We hypothesize that schooling dynamics are the result of positional and kinematic modulation of individuals responding to fluid dynamic stimuli from the movement of neighbouring individuals. We propose a two-tier approach to studying kinematic modulation within fish schools. First, quantification of the variation of individual movement in a school relative to that of a solitary individual uses an analytical pipeline combining artificial-intelligence-enabled tracking and video processing. Second, the study of kinematic modulation in response to hydrodynamic stimuli uses a mechanical flapping mechanism coupled with an enclosure to control fish position. We discovered that fish in schools exhibit higher levels of positional and kinematic modulation than individuals swimming alone. Fish swimming in enclosures can robustly respond to fluid stimuli from either a simple robotic fish or other fish located in proximity. This two-tier approach allows high-resolution analysis of positional and kinematic modulation within fish schools and their impacts on energy conservation resulting from collective movement.

## Introduction

A prominent idea in the study of animal collective behaviours is that the behaviour of the collective is the result of organized formations and synchronized movements of individuals within the collective. This introduces a conceptual dichotomy: if the collective is merely the multiplication of individual formations, why would the overall outcome of the collective be different from that of individuals moving alone? We hypothesize that the behaviour of the collective is, in fact, the result of individuals within the collective exhibiting different kinematic features from those of individuals moving alone. We predict that individuals moving within a collective should modulate their spatial movement and kinematics in response to their neighbours and exhibit different kinematic patterns than those shown when moving as solitary individuals. In the context of aquatic collective movement, *e.g.,* fish schools, this means that the individuals within a fish school modulate their kinematics to respond to fluid structures created by neighbouring animals and show different kinematics than when these same individuals swim alone.

Much of the existing literature presents fish schools as systems in which individuals show highly synchronized kinematics (Ashraf et al. 2016; Ashraf et al. 2017; Li et al. 2020; Mekdara et al. 2021) and move in organized formations in a planar configuration. Most of the analyses of the mechanisms by which fish schools experience benefits from collective movement focus on specific planar formations, where a few animals are treated as a functional unit (Weihs 1973; Thandiackal and Lauder 2023; Lauder et al. 2025). Although we acknowledge the value of focusing on formation swimming for models and theoretical analyses, the assumptions of synchronization, two-dimensional positioning, and fixed formations should be scrutinized when the results of models are extrapolated to the dynamic reality of three-dimensional biology.

Indeed, recent experimental observations have demonstrated that fish schools are a dynamic system where individual fish within schools constantly change their position and modulate their kinematics, and fixed planar formations are rare as fish frequently adopt three-dimensional relative positions (Ko et al. 2025).

Despite being a dynamic collective system, fish swimming in schools experience energetic savings relative to individuals swimming alone (Zhang and Lauder 2024; Zhang et al. 2024). This introduces an intriguing dichotomy: how can individual fish within schools exhibit dynamic interactions and not move in fixed relative planar formations while simultaneously saving energy? Resolving this dichotomy requires quantification of positions and body kinematics of individual fish within fish schools to examine the frequency distributions of kinematic parameters and compare them to those of the solitary individuals swimming alone. Previous studies estimate the links between the movement of individuals and locomotion energy saving using a kinematic analysis in a fish school on smaller time scales (Fish et al. 1991).

However, to understand variation in the position of individuals within a school and how individuals interact with each other, it is important to examine the variation of movement and kinematics within schools over a longer duration (Ko et al. 2025; Zhang et al. 2025).

We present a two-tiered framework for the investigation of collective dynamics of fish schools. Tier 1 focuses on analyses of the volume of schools, fish positions and kinematics in schools relative to individual fish swimming alone. Conceptually, the kinematic features of solitary individuals and fish in schools can vary in the breadth of a distribution curve as well as the locations and height of the peaks (Fig. 1A-C). We reason that these variations can be the result of the individual fish interacting with vortices generated by neighbouring animals (Fig. 1 D, E). Thus, we predict that individuals within a school should have greater variation in their positions and kinematic characteristics when compared with solitary individuals.

**Figure 1.**
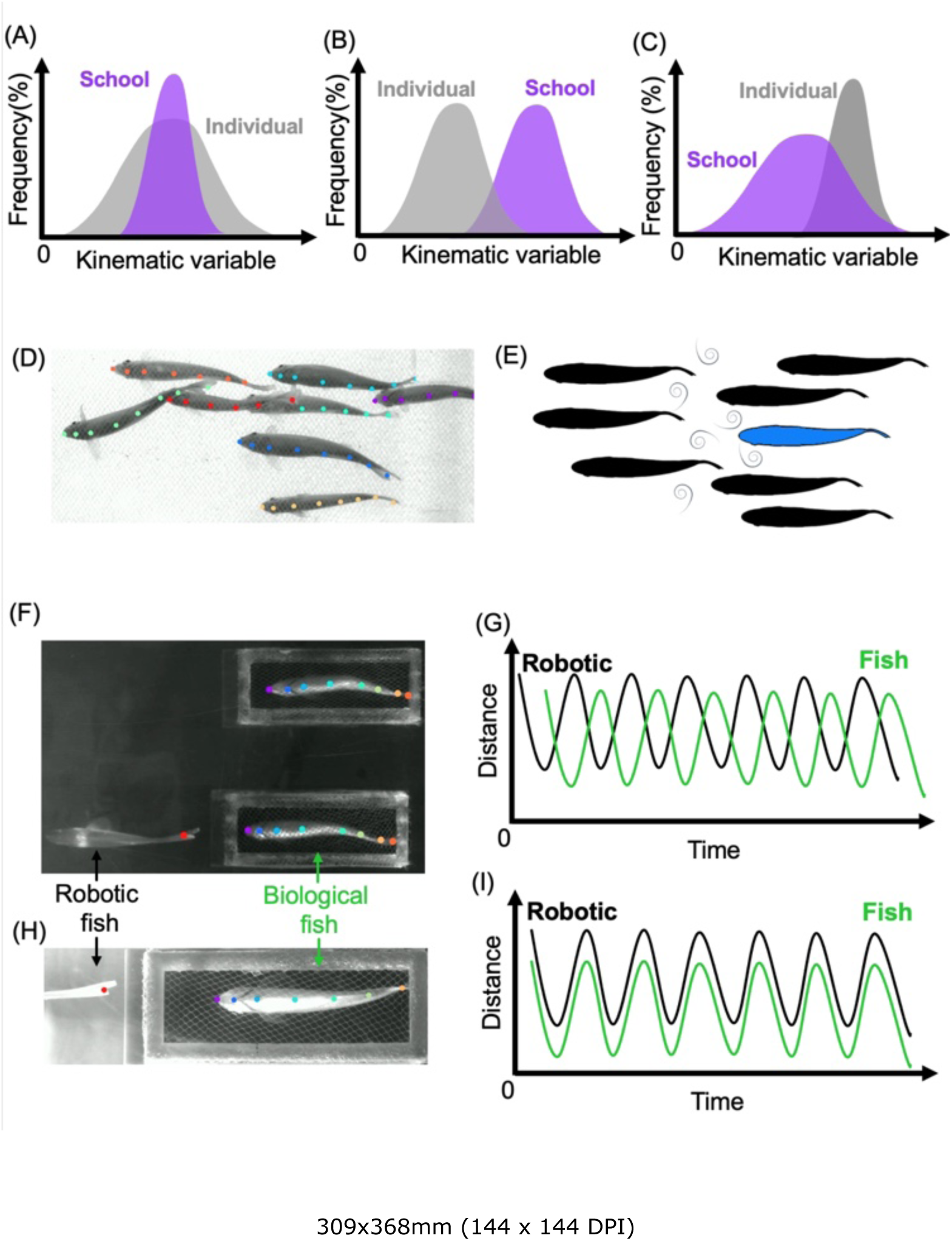
Conceptual framework of the two-tier approach showing (1) how the kinematics of individual fish can compare to those of schools, and (2) how individual fish can respond to the wake of neighbours. (A-C) Possible scenarios for variation of individual kinematics within a school compared to the kinematics of individuals swimming alone. Artificial-intelligence annotated frame (D) and schematics (E) show that the greater variation of individual kinematics within a school can result from individual responses to the vortex produced by neighbours. (F-I) Use of a robo-biological enclosure system to allow control over the relative positions of fish. This allows isolation of the kinematic responses of individual fish to the controlled vortex shedding frequency from either a robotic device or another fish in a cage upstream.

Our second approach, Tier 2, presents a robo-biological enclosure experimental system to study the kinematic and hydrodynamic effects of swimming in specific formations. The study of the effects of fish formation swimming has used primarily robotic and theoretical approaches (Kurt and Moored 2018; Kurt et al. 2021; Ormonde et al. 2024), although phalanx formations have been studied using fish swimming in shallow water systems to induce planar formations (Ashraf et al. 2017). We present this new experimental system as a guide to how the biomechanical analyses of collective movement can integrate a biological enclosure experimental system for live fish and a robo-mechanical system to provide a highly controlled experimental environment. We developed a platform where a small enclosure positions live fish near the wake produced by a robotic flapping mechanism (Fig. 1F-I). By varying the parameters of the flapping mechanism, we can mimic the vortex structures observed in fish schools and are able to repeatedly study the response of live fish to controlled variation in kinematic and hydrodynamic inputs.

## Materials and methods

### Model organism

Giant danio (*Devario aequipinnatus*), an active and schooling species, is our model organism. The animals were acquired from a local commercial supplier near Boston, Massachusetts (United States of America). Animals were randomly distributed and housed in separate schools in 37.9 l aquaria (*n* = 8 per tank). The aquaria have a self-contained filtration system, an aeration system (>95% air saturation, % sat.), and a thermal control (28 E&system. Fish were fed ad libitum daily (TetraMin, Germany), and 50% of the aquarium water was exchanged weekly. Animal holding and experimental procedures were approved by the Harvard Animal Care IACUC Committee (protocol number 20-03-3).

### Tier 1 experiments: high-speed videography

A fish school was tested in a rectangular (30 × 11.5 × 11 cm) carbon fibre enclosure (nylon netting) situated in a water tunnel that has a testing section of 50 × 30 × 30 cm. The use of the carbon fibre enclosure is to minimize the fish from seeking the boundary layer developed on the wall of the water tunnel (Ko et al. 2025). The nylon netting allows water to flow freely through the testing arena for recording the kinematics of fish school dynamics and ensures reduced boundary layer effects when fish swim near the enclosure sides. Three-dimensional schooling dynamics are being recorded using two streaming cameras (AOS PROMON U1000) fitted with a lens (AF-S Nikkor 17-35mm, Nikon) at 200 frames per second (FPS). The side-view and bottom-view cameras are 91 cm from the centre of the cage. The side-view was back-lit by an LED panel (HUION AS) that covered the entire field of view. The bottom-view was back-lit by two LED spotlights through diffusors (Nila Zaila). The shadows caused by the surface wave of the water tunnel were eliminated by a transparent plate placed on the water surface.

Eight giant danio were transferred into the carbon fibre enclosure the afternoon before each experiment, which was followed by a 15-h habituation period. During this time, the recording arena is covered by laser blackout sheets (Nylon Fabric with Polyurethane Coating) to prevent visual disturbance. Foot traffic to the experimental area was minimized to reduce audible disturbances. Altogether, these controlled testing conditions enable fish to exhibit their typical schooling dynamics and swimming behaviours in a laboratory setting. During the video recording period, water velocity in the water tunnel was incrementally increased from 0.5 to 8.0 body length per second (BL s^-1^) in increments of 0.5–2 BL s^-1^. Each increment lasted for 15 minutes, which enabled the fish to settle at each speed increment and exhibit typical swimming behaviours. For this analysis, we focused the analysis on video data collected at 6 BL s^-1^ because at this speed, schools of danio exhibit substantial energetic savings compared to individuals swimming alone (Zhang and Lauder 2024; Zhang et al. 2024). Throughout the experiment, fish were actively swimming against oncoming flow. The shrouding sheets and restricted foot traffic were also in effect during the experiment to reduce the effects of external stimuli.

### Tier 2 experiments: the robo-biological enclosure system

We designed and 3-D printed the enclosure system (35□30□78 mm, Fig. 4), a size that was specifically designed to accommodate giant danios. The support pillars of the enclosure used the shape of an airfoil (NACA 0012). Front and rear ends of the enclosure were grids with each component in a NACA 0012 shape, with the leading edge oriented towards the incoming flow. The top panel has an opening (20□43 mm) that serves as the means of inserting the fish into the enclosure. The side and bottom panels of the enclosures were covered in nylon netting that enables water flow, which is key for the fish to respond to the flow field introduced by the robotic system.

The robotic system was positioned in front of the enclosure cages and is a self-propelled flapping foil mechanical system (Lauder et al. 2007; Matthews et al. 2022; Thandiackal and Lauder 2023) attached to a 3-D printed fish manufactured in a flexible material (Formlabs 3+ printer; material: Elastic 50A). The robo-mechanical flapping mechanism is situated on a carriage suspended above the water tunnel. The flapper mechanical system is connected to a computer system that controls the location, pitching frequency and pitching angle. For the comparison of each relative formation, we include a control condition. The controlled condition is that the biological fish swim in the enclosure behind a static robotic fish.

### Particle image velocimetry for hydrodynamic analyses

To quantify the fluid dynamics of the robo-biological enclosure system, we used particle image velocimetry, as described in our previous research (Zhu et al. 2019; Zhang et al. 2024; Lucas et al. 2020). A horizontal plane of laser light was generated using a solid-state 532 nm green laser (LD solid-state green laser, 5 W, MGL-N-532A, Opto Engine LLC) that allowed visualization of flow through a central horizontal plane of the enclosure (Fig. 4). Flow velocity was calculated from sequential high-speed video frames using DaVis v8.3.1 (LaVision Inc., Göttingen, Germany). A vector field is generated by a sequential cross-correlation algorithm applied with an initial interrogation window size of 64 × 64 pixels that ended at 32× 32 pixels (overlap 50%, 2 passes at each window size). Particle image velocimetry was conducted to quantify flow patterns when the robotic system was flapping and generating fish-like wakes, but no fish were inside the enclosure. This is to quantify the effects of the enclosures on wake structures and assess if the fish inside the enclosure will encounter the fish-like wakes generated by the robotic system.

### Artificial intelligence-based tracking

We used both DLTdv8 (Hedrick 2008) and DeepLabCut (Mathis et al. 2018) for artificial intelligence (AI)-based tracking for kinematic analysis. We used the AI tracking functions in the DLTdv8 to track the tail movement of an individual fish within the enclosure.

AI-based tracking in DeepLabCut provided automated and robust tracking for two scenarios: 1) Kinematic responses of fish within the enclosure to the robotic fish; 2) Movement tracks and kinematic characteristics of the individuals within a fish school. Both the tracking of robotic fish and biological fish and the tracking of all individuals within the school used the multi-animal tracking function in DeepLabCut (Nath et al. 2019; Mathis et al. 2018). We tracked 8 markers on the fish body from snout to fork of the tail, and one marker on the tail fork of the robotic fish. We used a bottom-up neural network and manually annotated multiple frames, which were then used as a training dataset to train the neural network. The tracked videos were visually inspected for incorrectly tracked frames. We then re-labelled some of the incorrectly tracked frames and added them to the training dataset to retrain the neural network. We repeated the process until the neural network was able to track accurately.

### Automated image-based convex hull volume analysis

To continuously quantify the three-dimensional (3-D) occupancy space of the fish school, we estimated a 3-D convex hull volume using the 2-D convex envelope area for each view over each frame of the video (200 fps) in MATLAB (MathWorks, Inc.). The processing codes for 2-D convex envelope area used portions of codes that were modified from those of Ko et al. (2025).

We first extracted each frame’s image using the readFrame function and averaged over 1000 frames to estimate the horizontal and ventral background. We subtracted the background from the image of each frame to isolate the fish school. For each frame, we then set the image’s threshold to be –0.4 to scale with the darkness of the background and then removed white regions that are smaller than 500 pixels using the bwareaopen function. The bottom and front portions of the image were also converted to black colour by setting pixel values to 0. We then generated the horizontal and ventral 2-D convex area image for each frame of the bottom and side views, respectively, using the bwconvhull function.

The horizontal and ventral convex envelope areas were then used to estimate the 3-D convex hull volume. We isolated and converted the area of the convex envelope over each frame for both views from image format to Cartesian coordinates by finding the planar location of each pixel within the areas in the image using the find function. The ventral convex envelope area was then rotated to align with the long axis of the horizontal convex envelope area using the centroid of the two areas and the orthonormal eigenvectors of the horizontal convex envelope area extracted using Principal Component Analysis (PCA) (Mackiewicz and Ratajczak 1993). We then combined the horizontal and ventral convex envelope areas to generate the convex hull volume and calculated the three-dimensional schooling volume for each frame (Fig. 2A) using the convexhulln function, which is based on Quickhull (Barber et al. 1996). We averaged the schooling volume every 200 frames to generate the mean value and the standard error of the mean (s.e.m.) in MATLAB.

**Figure 2.**
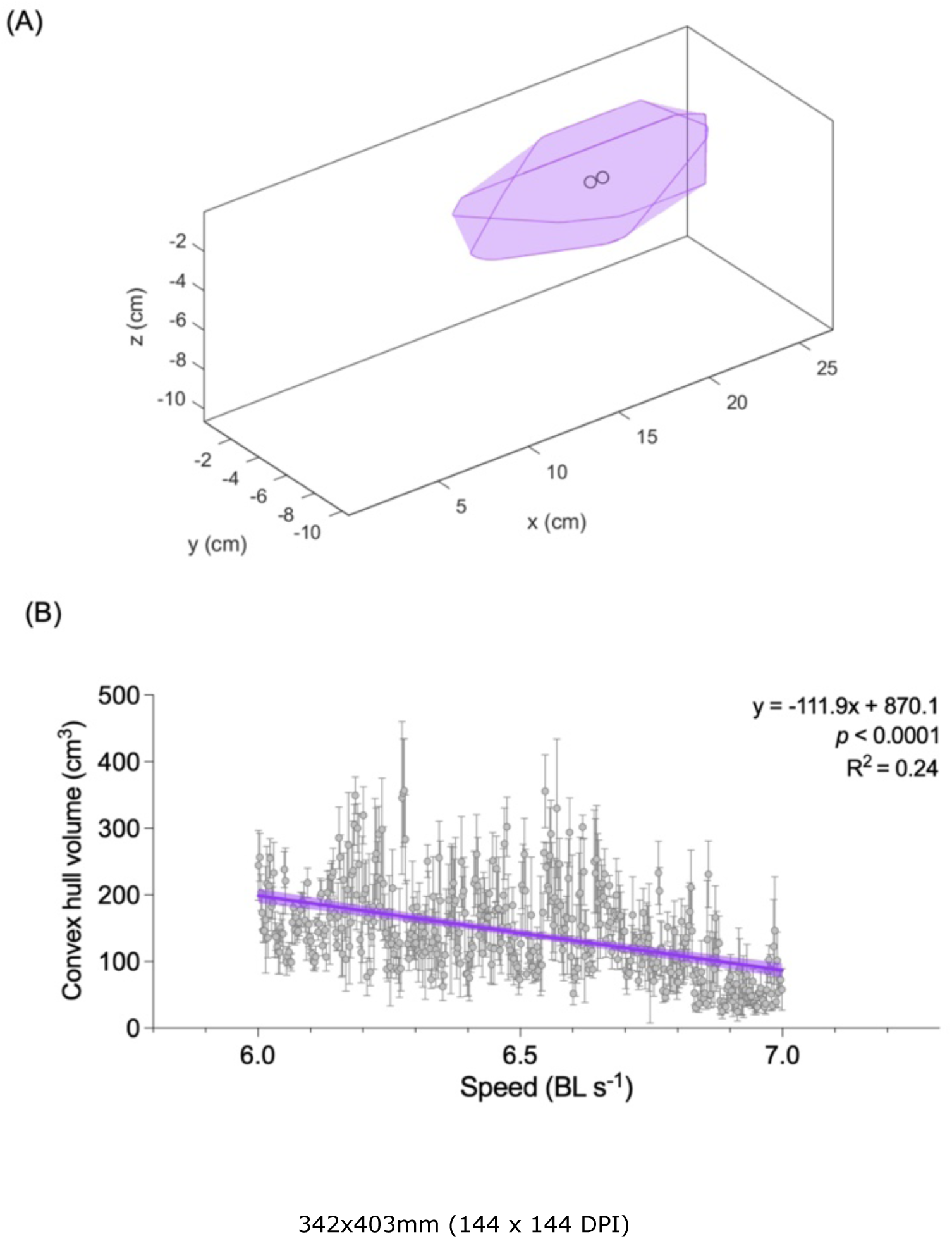
Three-dimensional analysis of school volume. (A) Example of the estimated three-dimensional volume of a giant danio (*Devario aequipinnatus*) school of eight individuals at one moment in time using a convex hull volume calculation. Circle corresponds to the centroid of the convex envelope (B) The three-dimensional volume of a fish school declines as flow velocity increases. The analysis used 15-min videos (200 frames per second) at each speed recorded in both ventral and lateral views.

To account for the effect of time for fish swimming within a 15-min speed increment at 6 BL s^-1^, we corrected the speed as the proportional duration the fish school spend in the increment (Brett et al. 1958; Brett 1964). A linear regression was fitted to describe the relationship between convex-hull schooling volume and swimming speed.

### Kinematic analysis

We quantified fish movement in the horizontal and vertical planes using tracked markers on the snout (Fig. 3A–D) to compare among individuals in the school and an individual swimming alone. We estimated the movement rate *v* of the snout of each individual fish over a window size *w* in the horizontal and sagittal view using bottom and side view video, respectively:

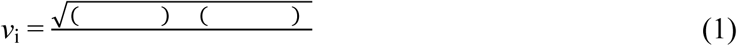

where *i* is the frame number, *x* and *y* are the tracked forward and lateral positions of the fish snout in each view, and *t* is time. We chose *w* = 150 frames as the window size for the movement rate *v* in this study.

**Figure 3.**
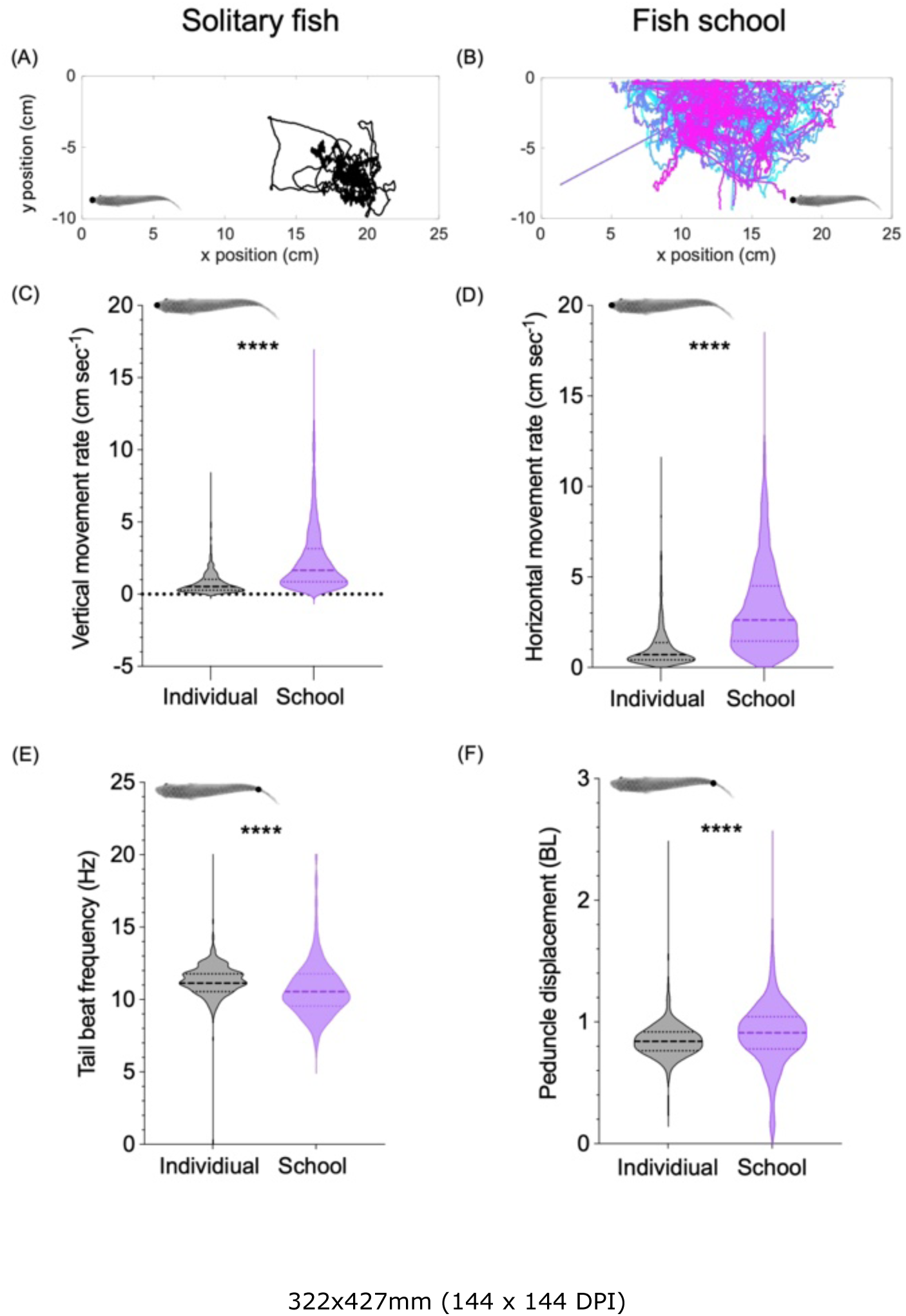
Positional and kinematic modulation of fish within a school in comparison to fish swimming alone. (A, B) 2-D movement tracks (ventral view) of individual fish swimming within a school and a fish swimming alone. (C, D). Violin plots of the horizontal and vertical movement rate for individual fish swimming in a school and a solitary individual swimming alone. (E, F) Violin plots of tail beat frequency and peduncle displacement for individual fish swimming in a school and a solitary individual swimming alone. The dotted lines are quartiles, and the dashed lines are the median. Data are from fish swimming at 6 body lengths per second (BL s^-1^).

We quantified body kinematics (tail beat frequency and peduncle displacement) using the position of the peduncle (Fig. 3E–F) in the horizontal view. We compared kinematics among individuals in the school and an individual swimming alone. To estimate the tail beat frequency we first identified the time when the maximal peduncle’s lateral position occurs. We then used the inverse of the time difference between consecutive maximal positions to calculate tail-beat frequency. To estimate peduncle displacement, we first identified consecutive pairs of the maximum (peak) and minimum (valley) values in a cycle of the peduncle’s position. We then calculated the difference between the maximum and minimum values at the horizontal plane for each tail beat cycle to estimate peduncle displacement.

### Statistical analyses

Welch’s t-tests were used to compare the average values of kinematics metrics between individuals swimming alone and fish within the school. A linear mixed-effect model (speed and locations of the flow) was used to analyze the effects of enclosure on the velocity field (*post hoc* test: Two-stage Benjamini, Krieger, & Yekutieli FDR procedure). The effect of vortex shedding from the robotic flapping mechanism on the in-line and staggered positions of the enclosure with fish inside was studied using one-way ANOVA (*post hoc* test: Holm-Šídák). The analysis of fish kinematics staggered behind the robotic fish used Mann-Whitney tests. All statistical analyses were conducted in Prism (v. 10.6). The spectral analysis of kinematic trade-offs was conducted using the power spectral density function using tail beat frequency and tail beat amplitude using ADInstruments LabChart (v. 8.1).

## Results

We discovered that individuals in a school had higher levels of positional and body kinematic variation than solitary individuals swimming alone. Biological fish in enclosures responded to fluid stimuli from a neighbouring robotic fish by altering their swimming kinematics.

### Individuals within fish schools have higher kinematic variation than fish swimming alone

Although the three-dimensional volume of schools declined as swimming speed increased (y = – 111.9x + 870, where y is the schooling volume, and x is swimming speed; *p* < 0.0001; R^2^ = 0.24) (Fig. 2), individual fish within the school traverse nearly two times larger area than fish swimming alone (Fig. 3A, B) at 6 BL s^-1^ flow velocity. The larger movement areas of fish within the school corresponded to the significantly higher movement rates in both horizontal and sagittal planes (movement rate: horizontal plane = 3.3 cm s^-1^; vertical plane = 2.4 cm s^-1^) than those of fish swimming alone (movement rate: horizontal plane = 1.2 cm s^-1^; vertical plane = 0.84 cm s^-1^; *p* < 0.0001; Fig. 3C, D). Despite the larger rate of movement for fish within the school, these fish had a lower tail beat frequency (10.7 Hz) than that of the fish swimming alone (11.3 Hz; *p* < 0.0001; Fig. 3E), which was accompanied by larger peduncle displacement (0.9 BL) than that of the fish swimming alone (0.84 BL; *p* < 0.0001; Fig. 3F).

### Hydrodynamic effects of the enclosure

The enclosure has small yet detectable effects in reducing fluid velocity (Fig. 4). Freestream flow velocity and the locations of the flow field relative to the enclosure have a significant interaction (*F*_6,32_=18.8; *p* < 0.0001). We use the flow field in front of the enclosure as the control condition. The effects of enclosure are demonstrated by comparing the flow field in the centre and rear of the cage to that at the front of the enclosure. Across the tested speed range, the centre of the enclosure, where fish would locate, experienced an average streamwise flow velocity reduction from 2.3 to 4.0% (*p□*0.029, *F*_2,32_ = 200.4) when compared to the flow field at the front of the cage. A reduction of average downstream flow velocity from 7.7 to 11.3% across the tested speed range existed at the rear of the cage (*p* < 0.001, *F*_2,32_ = 200.4) when compared to the flow field at the front of the cage (Fig. 4). Although the enclosure impacted the hydrodynamic flow field, the oscillatory accelerated wake generated by the robotic fish anterior to the cage was still present (Fig. 4).

**Figure 4.**
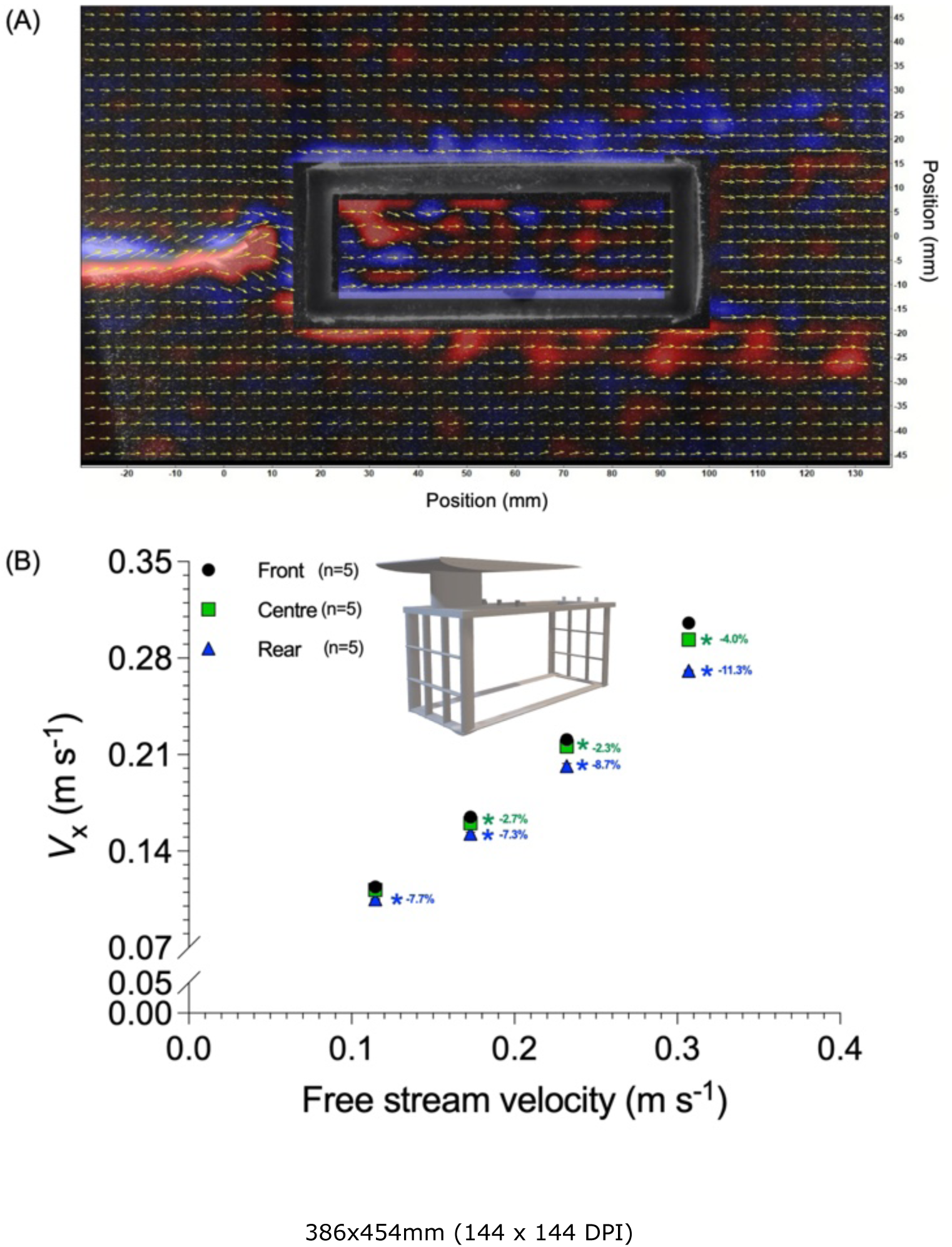
Robo-biological experimental platform facilitating a controlled analysis of fish relative positions and hydrodynamic and kinematic effects. (A) Testing for the hydrodynamic impact of the enclosure system on the reverse Kármán vortex street generated by a leading robotic fish. Wake features are retained and pass through the enclosure system. Testing is done under laminar flow conditions. (B) Measurement of the flow velocity in front of, inside, and behind the enclosure. The fluid drag induced by the enclosure is not more than 4% of the freestream flow.

### Kinematic responses of fish within an enclosure

We programmed a flexible fish model to swim in front of the enclosure containing a biological fish to assess the extent to which the fish would alter kinematics in response to an oncoming flow pattern (Fig. 5A, B). The biological fish in the enclosure exhibited steady-state undulatory swimming to match freestream velocity (20.7 cm s^-1^) and exhibited a tail beat frequency of 2.1 0.015 Hz. Once the robotic fish started undulating (2 Hz, 10-degree pitch amplitude), the biological fish began to modulate body kinematics to match the robotic flapping frequency (Fig. 5C). Within approximately three tail beat cycles, the biological fish reduced its tail beat frequency from 4.8□0.045 Hz to 2.1 Hz, matching that of the robotic fish for seven tail beat cycles (Fig. 5C). Although the tail beat amplitude of the biological fish increased from 4.1 to 5.7 mm at the testing speed, the difference was not statistically significant (*F*_2,51_ = 5.23; *p* = 0.18; Fig. 6). The biological fish synchronized tail beats with the robotic fish at exactly 2 Hz for five tail beat cycles (Fig. 5C): tail beat frequency synchronization of biological and robotic fish demonstrated that fish within the enclosure can respond to the vortex wake shed by the undulatory motion of an anterior in-line flapping system. Frequency synchronization was also observed between the tail beat frequency of the robotic fish and the frequency of mid-line undulation of the biological fish (Fig. 5D). Moreover, compared with the control condition, the biological fish responded to the flapping robotic fish by displaying a trade-off in the reduced tail beat frequency and a more variable (almost bimodal distribution) tail beat amplitude (Fig. 5E).

**Figure 5.**
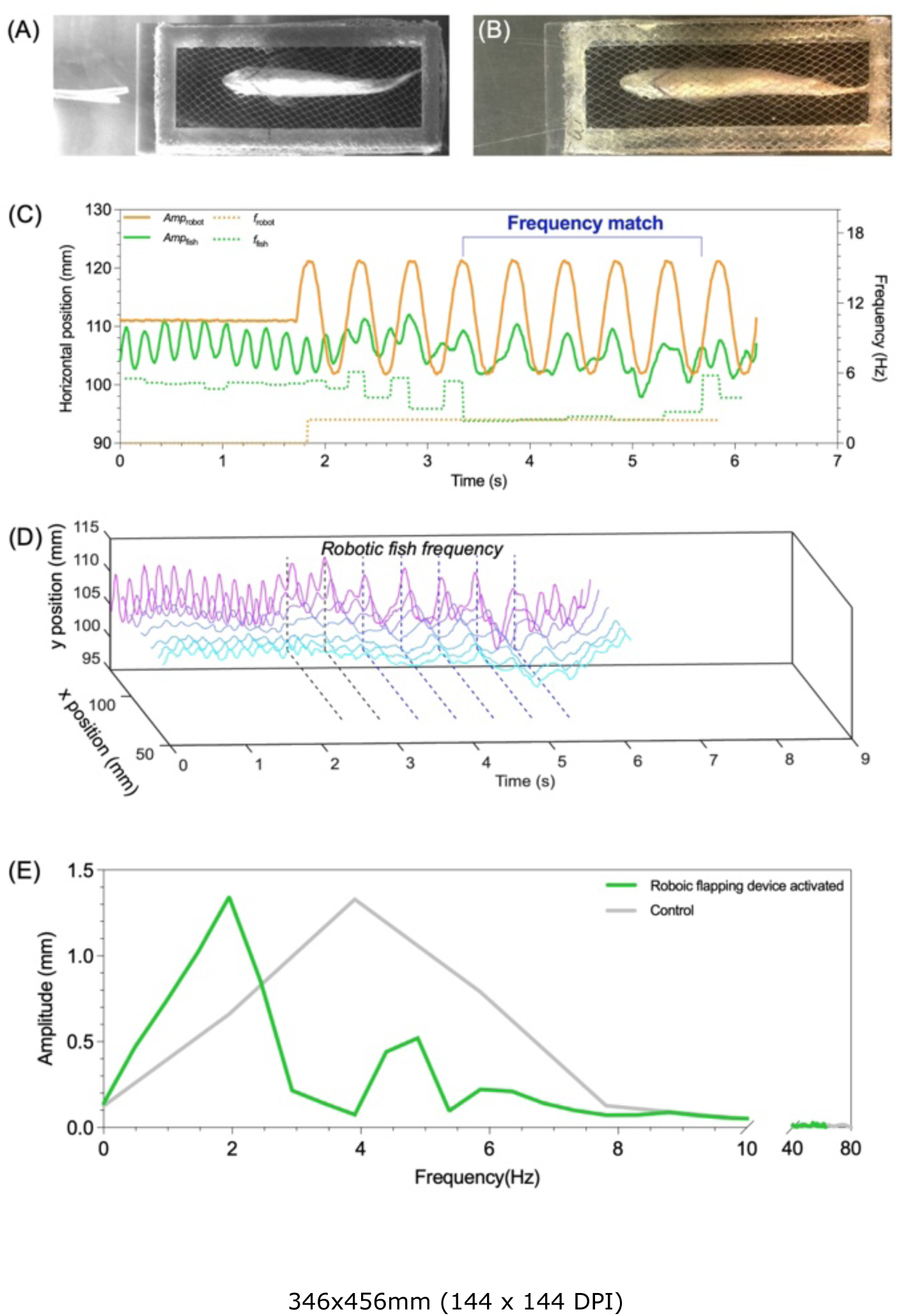
The response of fish to the wake produced by a leading robotic fish. (A, B) Fish inside the enclosure behind a robotic fish (A) and without the robotic fish (B). (C) Frequency synchronization of the tail beat between the fish and the robotic fish. (D) Position of different locations along the mid-line (denoted by different colored lines) over time, showing oscillatory frequency of the mid-line of the fish) matching the oscillatory frequency of the robotic fish (dashed line). The black dashed lines annotated the robotic and biological fish undulatory frequency was approaching synchronization. The blue dashed lines annotated the undulatory frequency are synchronized between biological and robotic fish. (E) Spectral analysis of the shift in tail beat frequency and tail beat amplitude of the fish in response to the oscillatory motion of the robotic fish. The freestream flow velocity was 20.7 cm s^-1^.

**Figure 6.**
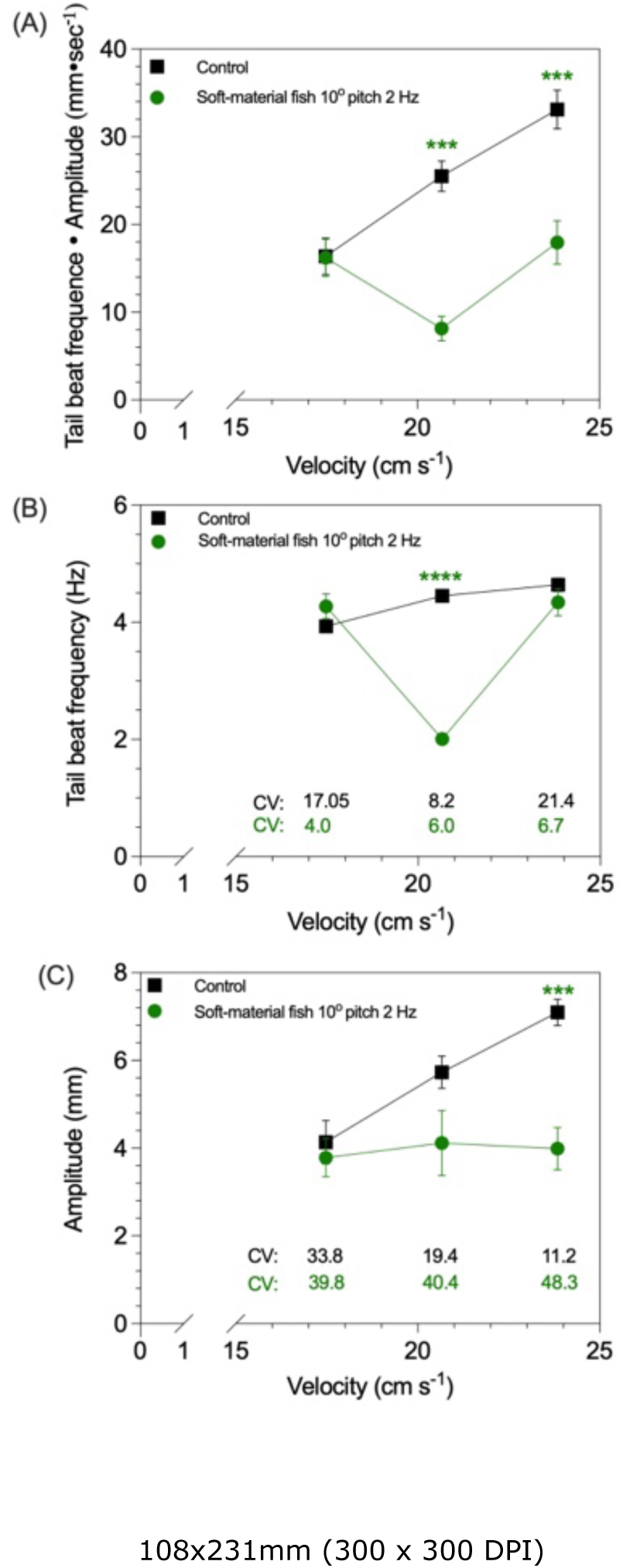
Kinematic analysis of a fish within an enclosure swimming in-line behind a robotic fish. (A)Total kinematic effort (Tail beat frequency□Amplitude), (B) tail beat frequency, and (C) tail beat amplitude of the fish swimming in the enclosure behind a robotic fish oscillating at 2 Hz and an amplitude of 10. Coefficients of variation (CV) are provided for the tail beat frequency and amplitude measurements. Asterisks denote the statistical significance (\*\*\**p* 7T0.001, **** *p* 7T0.0001).

We varied testing speed to understand how the kinematics of biological fish respond to the momentum wakes generated by undulating robotic fish as a function of speed. Compared with the control condition of a biological fish swimming in the enclosure located in freestream flow, a biological fish swimming in the enclosure behind a robotic fish showed reduced kinematic effort (measured as tail beat frequency times amplitude) at flow a velocity of 20.7 & 23.8 cm s^-1^ (*p□*0.0003, *F*_2,51_ = 9.04), but not at 17.5 cm s^-1^ (Fig. 6A). Moreover, the contributions of amplitude and frequency to the reduction in kinematic effort differed between medium and high speeds. The reduced kinematic effort at the medium speed was achieved by a lower tail beat frequency (*p* < 0.0001, *F*_1,23_ = 12.5) (Fig. 6B), whereas the reduced kinematic effort at the high speed was achieved by a lower tail beat amplitude (*p* = 0.0001, *F*_2,51_ = 5.24) (Fig. 6C). Furthermore, biological fish had a lower variation in tail beat frequency but a larger variation in tail beat amplitude when following momentum wake of the robotic fish (*see* values of coefficient of variation).

When compared to undulatory motion in the freestream, biological fish located laterally to the robotic fish moved at 4 Hz with 10 amplitude in the staggered position, had a higher tail beat frequency (*p* = 0.0295; Fig. 7). However, we did not observe a statistically significant difference of tail beat amplitude and the kinematic effort of the biological fish that was staggered behind the robotic fish when compared to the control group (*p□*0.311; Fig. 7).

**Figure 7.**
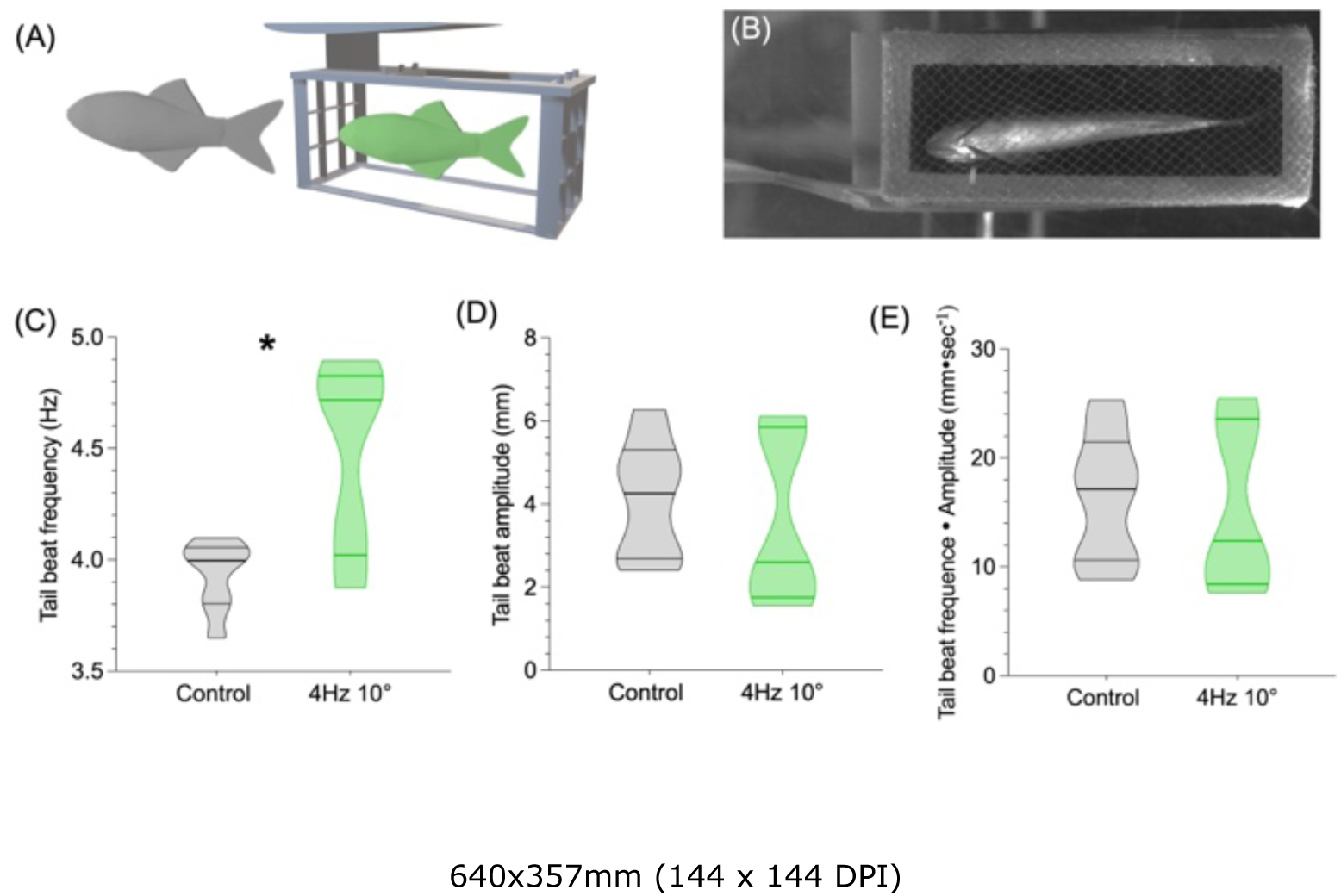
Kinematic analysis of a fish within an enclosure swimming in a laterally staggered behind a robotic fish. (A, B) 3-D model illustration and a picture of the relative position of a fish within an enclosure that is located in a staggered position lateral to a robotic fish. (C) Tail beat frequency, (D) tail beat amplitude, and (E) total kinematic effort (Tail beat frequency□Amplitude), of a fish swimming laterally staggered behind a robotic fish. The robotic fish was oscillated at 4 Hz and an amplitude of 10. The freestream velocity was 17.3 cm s^-1^. Asterisks denote the statistical significance (*, *p* 7T0.05). The horizontal bars in the violin plots are the 25 and 75 percentiles and the median.

## Discussion

The two-tier approach to studying fish schooling dynamics suggested here aims to both improve our understanding of the overall dynamics of fish schools and to also allow study of fluid dynamic and kinematic interactions, as well as the effects when individuals within a fish school maintain, even for a short time, relatively stable positions (Fig. 1). The first-tier approach aims to characterize fish schooling dynamics over a long duration (hours) and in three dimensions, which is achieved through an artificial-intelligence (AI) based tracking and image-based analytical pipeline. By using a smaller enclosure with net walls located within a larger flume, boundary layer effects are substantially reduced, and three-dimensional data on fish schooling behaviour can be obtained (Ko et al., 2025). Although artificial intelligence generates an abundance of data, the challenge is to extract meaningful biological understanding. Using fish schools as a model system, the proposed analytical approach shows the potential effectiveness of AI-based tracking for studying the biomechanics of collective movement over longer time frames at a high frame rate.

There can be relatively short instances when fish maintain somewhat stable positions relative to each other within a school (Ashraf et al. 2017; Ashraf et al. 2016; Ko et al. 2025; Lauder et al. 2025). These brief interactions provide insight into the mechanisms of how kinematics interact with the specific fluid structures; however, the challenge is that these interactions are transient and difficult to consistently reproduce. Hence, second tier approach focuses on studying the kinematic and fluid dynamic interactions among individuals using an experimental arrangement where individual fish can be subjected to a highly controlled and repeatable stimulus. We propose using a novel robo-biological enclosure system to both control the position of the live fish and the local fluid field that the live fish will experience. Using a mechanical flapping system with a flexible model fish, we can generate known wake patterns of specific frequency and amplitude and subject the fish within the enclosure to these stimuli. To account for the effects of the enclosure, we matched the treatment groups with control groups where the same testing conditions were maintained, and the only variable was the presence or absence of the robotic stimulus. Collectively, the two-tier approach enables the analysis of positional and kinematic modulation within fish schools and their potential impacts on the energy conservation resulting from collective movement (Zhang and Lauder 2024; Zhang and Lauder 2025; Nguyen and Peleg 2025).

### Tier 1: dynamic fish schooling structure and kinematics

The decrease of three-dimensional schooling volume with speed (Fig. 2), which is quantified at a a high resolution of 200 counts per second, agrees with previous studies (estimated at a lower resolution) that schooling volume shrinks as speed increases (Zhang and Lauder 2024; Zhang et al. 2024; Ko et al. 2024). This generally supports the notion proposed by the previous study that fish schools as a whole act as a shelter for the individual fish within when hydrodynamic conditions become challenging (Zhang et al. 2024). The high-resolution quantification, however, reveals that the schooling volume is highly dynamic and shows large variation even within a 15 min period (coefficient of variation = 46.6%).

We have discovered that individuals within the fish school have a higher rate and extent of spatial movement when compared to individuals swimming alone (Fig. 3). Many studies of fish schooling behaviour have failed to make comparisons to a control condition in which fish school dynamics are compared to the pattern of movement by individuals swimming alone at the same speed. This comparison (Fig. 3) has revealed that solitary locomotion is *less* variable than when individuals move within a school. Intriguingly, the higher movement rates were not driven by an increase in tail beat frequency. Instead, the peduncle displacement amplitude of fish within the school increased. The decreased frequency of fish within a school with increased amplitude is an indirect suggestion that there may be a reduction in energetic cost. Swimming with increased amplitude can result from a reduction in body muscle activation as fish take advantage of oncoming flow, and suggests similarity with the Kármán gait, a feature of passive undulatory motion in which energy expenditure is considerably reduced (Liao et al. 2003a; Liao et al.

2003b; Beal et al. 2006; Taguchi and Liao 2011). Previous studies of this same giant danio species have demonstrated that, at the swimming speeds tested here, individual fish in schools reduce locomotor energy use relative to solitary locomotion (Zhang and Lauder 2024). Future studies are needed to understand the potential mechanisms that underlie positional and kinematic modulation and how these are connected to energy saving by individuals within schools.

The statistical characterization of schooling dynamics is best achieved by analyzing large datasets generated by artificial intelligence-enabled tracking. The field of collective behaviour broadly encompasses terrestrial, aerial and aquatic organisms (Bazazi et al. 2008; Portugal et al. 2014; Strandburg-Peshkin et al. 2015; Ling et al. 2019; Coulombe et al. 2025), and over the years, studies have moved from marker-based tracking to marker-less tracking, leveraging artificial intelligence (Crall et al. 2015; Delacoux and Kano 2024; Bozek et al. 2021; Smith et al. 2022). Unlike other organisms, attaching markers to fish comes with its own set of challenges, *e.g.,* the mucus excretion and marker retention on the flexible body (Lauder et al. in press).

Although fish can be marked with visible implant elastomer tags (Curtis 2006; Jungwirth et al. 2019), the tags can affect social preferences as has been observed in zebrafish (Frommen et al. 2015). Marker-less tracking can result in mistracking of individual identities where manual corrections are needed. Tracking multiple landmarks on each individual’s body within a collective continuously over a long duration and studying the related biomechanics and fluid dynamics, as proposed in the two-tier approach, has not often been accomplished. This is due in part to the challenges of individual identity tracking when overlaps occur, which might be overcome by AI tracking models rooted in the three-dimensional space that resolves occlusion.

### Tier 2: kinematic and hydrodynamic interactions during in-line swimming

In-line swimming is an intriguing formation that has been shown to have the second-highest presence (14%) among four documented formations within three-dimensional fish schools (Ko et al. 2025). A counterintuitive aspect of fish in-line formations is that the region right behind a fish’s caudal fin can generate a thrust wake with increased velocity that increases the drag for the immediate follower. Hence, the follower might not be able to obtain hydrodynamic benefits for energy saving. Previous research using a robotic model (Saadat et al. 2021) has suggested that, despite this potential drawback, energy saving by in-line swimming is possible. In addition, a study using a flapping foil as a model for the upstream fish of a tandem pair (Thandiackal and Lauder 2023) showed that a trout swimming behind a flapping foil was able to synchronize tail beat frequency with the flapping airfoil. Hydrodynamic analysis demonstrated that the follower obtains thrust through both the oscillating flow and through modification of vorticity that passes along the body and through head oscillation that enhances leading edge suction on the follower (Guo et al. 2025). Benefits for the following fish derive in part from the Knoller-Betz effect of leading-edge suction, which occurs when the oscillating wake from a leading airfoil-shape objects result in lift and thrust on the head of the following fish (Knoller 1909; Betz 1912; Akhtar et al. 2007).

However, these previous experiments used an airfoil that is much larger than the depth of the following fish, and thus do not represent the actual schooling case of two fish in series. Since the wake produced by the airfoil is much larger in depth when compared to the body depth of the trout (Thandiackal and Lauder 2023), the trout was swimming in a simplified 2-D flow structure rather than a 3-D flow structure that results from the wake behind a leading fish. By using a flexible model fish that is the same size as the giant danio within the enclosure (Fig. 4), we were able to test the responses of live fish to more biologically relevant flow structures present within a school. The results show that followers in the in-line formation respond to the more realistic 3-D flow structure that more closely resembles that of the wake generated by fish (Fig. 5). The follower synchronized tail beat frequency with that of the robotic fish and demonstrated a reduced tail beat frequency and increased tail beat amplitude, similar to the kinematic features of the Kármán gait that conserves locomotor energy (Liao et al. 2003a; Liao et al. 2003b; Beal et al. 2006).

### Kinematic and hydrodynamic interactions of a staggered formation

Often, a following fish might not swim perfectly in line with a leading fish. Hence, we want to explore kinematic responses when a following fish is horizontally staggered behind a leading fish with an offset in mean swimming trajectory using the robo-biological enclosure system (Fig. 4). Staggered formations have been studied previously from a computational fluid dynamic perspective using data from schooling fishes (Lauder et al. 2025; Huang et al. 2025; Guo and Dong 2025) and have been shown to be advantageous in reducing energy consumption during swimming (Huang et al. 2025) (Kurt et al. 2020). The horizontally staggered formation can be viewed as a sub-section of the classic “diamond formation” where each fish in the formation is horizontally staggered off each neighbour (Weihs, 1973). In the horizontal plane, the staggered configuration is shown to be a common formation at low swimming speeds (e.g., ∼0.8 BL s^-1^), while a phalanx configuration becomes more dominant at higher speeds under conditions where fish are confined to relatively shallow water, in a planar configuration (Ashraf et al. 2017). A staggered formation can occur in both the horizontal and vertical dimensions (Ko et al. 2023; Menzer et al. 2025). Long-duration kinematic analyses suggest that the vertically staggered formation (ladder formation, present for 79% of the time) is more abundant than any other formation, whereas the horizontally staggered formation occurs only 7.6% of the time (Ko et al. 2025). Our experimental data using a robo-biological enclosure system (Fig. 7) show that fish horizontally staggered behind a flapping robotic mechanism appear to increase tail beat frequency, while other kinematic metrics, such as kinematic effort, remained the same. Analyses of staggered formation swimming using live fish to assess the ability of each fish in the pair to alter their kinematic pattern have yet to be conducted over a broad range of frequency and amplitude parameters. Such future studies promise to contribute to the role that staggered formations play in schooling dynamics. The robo-biological platform provides one controlled experimental approach to understanding the broad category of laterally staggered fish planar formations.

### Use of a robo-biological experimental platform

The use of robotic systems to study animal biomechanics has contributed to understanding how animals modulate their kinematics in response to changes in the aerial, terrestrial and aquatic environments (Gravish and Lauder 2018; Ramdya and Ijspeert 2023; Ishida et al. 2024). The advantage of the robotic systems is the ability to systematically sweep the parameter space and to test a specific hypothesis under highly controlled conditions at the intersection of physics and physiology (Lauder 2022). Often, such experimental control is impossible to achieve in studies of live animals alone, where controlling behaviour can be challenging. Here, we introduce a robo-biological experimental platform as a hypothesis-probing device to test specific biomechanical mechanisms at the level of the individual, and understand how these scale up to the level of animal collectives. The main benefits of such a controlled approach to understanding the interactions of two (or more) individuals are (1) the ability to generate a reproducible fluid field that mimics the ones generated by neighbouring individual fish, and (2) the capacity to repeatedly measure kinematic features of live animals when interacting with the fluid field.

The robo-biological experimental platform is not without its limitations. The tight spatial constraints means that fish cannot track the movement of the vortex outside the enclosure. As we demonstrated in the Tier 1 approach, the individuals within the school exhibited an abundant number of movements in relation to each other. Although the soft-body robotic fish produced wakes that are closer to the biological fish than a 2D airfoil, the real fish often do not undulate at the set frequency and amplitude and at the same location. Hence, the robotic fish produces a more consistent fluid stimulus than that of the biological fish. Instead, a real fish leading another moves laterally and vertically, which prompts the follower fish to respond and potentially coordinate movement. These dynamic interactions cannot yet be replicated in the robo-biological experimental platform. However, future iterations can potentially include such dynamic movements and simulate the more realistic inputs to a real following fish.

### Conclusions

The study of the collective movement of fishes (or other model organisms) is a complex problem that lies at the intersection of physics and physiology (Zhang and Lauder 2023; Zhang and Lauder 2025; Nguyen and Peleg 2025). Research in this field is also currently leveraging cutting-edge developments in technologies such as robotic systems, computational fluid dynamics, and AI-enabled tracking (Hedrick 2008; Lauer et al. 2022; Guo and Dong 2025; Ormonde et al. 2024; Kelly et al. 2023). One of the most significant challenges in studying fish schooling behaviour, biomechanics, and the hydrodynamic mechanisms that mediate inter-individual interactions in a school, is that the collective movement of fish schools is a highly dynamic system where group movement in fixed relative positions does not always occur.

Indeed, fish swimming in schools can show increased variation in kinematics and positioning relative to individuals swimming alone, nearly constantly repositioning and changing their formation. This begs the question of how the fish school as a dynamic collective can conserve locomotor energy when the fish within are repositioning and not consistently maintaining specific formations.

To address this question, we propose studying fish schooling dynamics using a robo-biological experimental platform to reproduce the hydrodynamics and kinematic responses of inter-individual interactions (Lauder 2022). Controllability and reproducibility are the advantages of robotic and robo-biological systems compared to when two variable biological systems interact (Li et al. 2024; Roderick et al. 2021; Liu et al. 2026; Wang et al. 2023; Gravish and Lauder 2018; Lauder 2022). Acquiring long-term three-dimensional kinematic data on fish schools to allow analyses of kinematic modulation by individuals within the school, coupled with the use of a robo-biological platform to study specific interactions, is poised to provide a research path forward for studying the complex biology of collective movement.

### Future Directions

The study of schooling biomechanics is at the cusp of making the leap from individual kinematics to collective dynamics (Lauder et al. 2025). In order to understand the physics and physiology of schooling behaviour in fishes (Zhang and Lauder 2025; Nguyen and Peleg 2025), we must move beyond studies of fish groups moving in still water to analyze how school structure changes as swimming speed increases, and to compare any such changes to those displayed by individual fish moving alone over the same range of speeds. The complexity of fish schooling behaviour requires that we use some approach to manipulate the school, and changing schooling speed using a flow tank is one key experimental manipulation that can help reveal inter-individual schooling dynamics. Ideally, in the near future, kinematic analysis should move towards three-dimensional space and use the artificial intelligence-based tracking models built for true three-dimensional analysis, which is still challenging at the present time.

## Author contributions

Conceptualization: YZ, GL

Methodology: YZ, DR, GL

Investigation: YZ, DR, GL

Visualization: YZ, DR

Funding acquisition: GL, YZ

Project administration: GL, YZ

Supervision: GL

Writing – original draft: YZ

Writing – review & editing: YZ, DR, GL

## Acknowledgements

Many thanks to members of Lauder Laboratory for numerous discussions about fish schooling behaviour, for comments on the manuscript, and to Cory Hahn for fish care. Thanks Hungtang Ko for sharing the base codes of imaging processing.

## Data and materials availability

All data are available in the main text.

## Figure Captions

**Supplementary Movie 1. An illustration the convex hull volume of a fish school changing with time**. The analyses were conducted for each frame for a short video recording at 200 frames per second.

